# Chronic infection alters pathogen virulence, microbiome composition, and fly physiology across generations

**DOI:** 10.1101/2024.03.04.583275

**Authors:** Krystal Maya-Maldonado, Nichole A. Broderick

## Abstract

In many insects, parents and offspring share the same environment. Thus, an infection in the parents has the potential to influence offspring defenses. Moreover, infection can also affect other host aspects, including the microbiome, development, and reproduction. To better understand the intergenerational impacts of infection, we assessed the effects of challenge by the gut pathogen *Pseudomonas entomophila (Pe)* on *Drosophila melanogaster*. We found that parental challenge by *Pe* led to environmental transmission of the pathogen from parents to offspring, resulting in a persistent infection among the population. *Pe* is a highly virulent pathogen; however, we found that persistent infection was correlated with a loss of pathogen virulence across generations. We explored the impact of chronic pathogen exposure on host physiological traits. Our results showed that pathogen load, virulence, and pathogen-induced microbiome remodeling influence fecundity and starvation resistance. Current research in *Drosophila* and other insects has shown that immune status can be transmitted to the next generation (transgenerational immunity). Since the offspring were continuously exposed to the pathogen, we explored their response to a new infection. Even though we did not find a protective effect, we observed alterations in gene expression and microbiome remodeling following a new *Pe* challenge that was dependent on the parental treatment. Altogether, our results provide evidence that the pathogen adapted across generations as part of a tolerance mechanism that allows the pathogen to persist in the environment, which confers a greater probability of survival in subsequent generations. However, chronic exposure to the pathogen resulted in a cost to the host by altering several aspects of host physiology.

**Author summary:** Infection impacts many aspects of animal physiology, including priming host immune responses to repeated pathogen exposure. Whether parental experiences with a pathogen can influence such responses in offspring is less certain. Here, to further our understanding of generational impacts of infection, we studied the interaction between host immunity, the microbiome, and a gut pathogen across generations using the model organism *Drosophila melanogaster.* Our results showed that parental challenge established a persistent infection in the population, such that offspring were chronically exposed to the pathogen. This chronic pathogen exposure impacted many host physiological traits, but did not confer protection to re-infection with a high-dose of the pathogen. Instead, we found that the transmitted infection led to a loss of pathogen virulence in offspring. At the same time, pathogen density, virulence, and pathogen-induced microbiome remodeling influenced fecundity and starvation resistance. Overall, our results highlight that infection in parents can influence intergenerational responses due to impacts both on the microbiome and on selection on pathogen virulence. Such chronic interactions with the pathogen, even reduced in virulence, alter host physiology.

## Introduction

The immune system is a crucial modulator of host-pathogen and host-microbiome interactions (1–3). Most studies on host immune responses to microbial challenge focus on impacts within the same generation. As such, host-pathogen and host-microbiome interactions across generations are relatively unexplored. The natural ecology of many insects permits the inheritance of microbial infections either by direct vertical transmission from parents to offspring, or by environmental acquisition (4–6). Such associations allow the spread of pathogens through populations, with the potential to cause persistent infections. Chronic exposure to a pathogen is largely thought to have negative effects on host physiology. However, prior exposure to a pathogen can also positively shape the immune response, and stored information from such previous encounters can be recalled to mount a protective response upon a new exposure to the pathogen (7). In insects, this response is called immune priming or innate immune memory and has been shown in some circumstances to impart protection even across generations, known as transgenerational immune priming (TGIP) (8–10). Several criteria, including the ecology of the host, infection procedure, pathogen load, host basal immune status, fitness costs of infection, and the role of the microbiome have been proposed as key aspects to consider when such response is investigated (9,10).

In all animals, including *Drosophila,* the innate immune system interacts intimately with the microbiome, which influences several aspects of host biology, including digestion, nutrition, host gene expression, morphology, and epithelial architecture of the gut (11–13). All these aspects are vital for proper gut homeostasis and their alterations may have effects on host defense. The microbiome maintains a basal expression of immune effectors, such as antimicrobial peptides (AMPs), through IMD pathway activation (11,14,15). Such host gene expression, regulated by the microbiome, has the potential to influence subsequent generations. Pathogen interactions within or across generations have the potential to induce host alterations beyond immune responses. As such, previous pathogen encounters may lead to trade-offs between immune benefits and negative fitness effects or vice versa, including metabolic, reproductive, and abiotic stress in several insect species (16–20). In *Drosophila*, the microbiome can affect such physiological alterations across generations (21,22). However, innate immune memory within or across generations has rarely been examined in the context of the normal host microbiome.

At the same time, in nature, insects often grow, feed, and reproduce in a limited niche, meaning they can share several factors that can influence biological processes as a community, even when they are not eusocial insects. *Drosophila* exhibit social behaviors including group foraging and gregarious oviposition, and as such adult flies influence the food and environment in which offspring develop (fermenting/rotting fruit in nature or a food vial in the laboratory) (23,24). When parents and offspring share the same environment, the offspring are more likely to be influenced by parental experiences, including encounters with pathogens. Untangling such influences is complex, and *Drosophila* provides a good model to consider several pieces of the puzzle: parental experience, offspring impacts, and the role of commensal and pathogenic bacteria.

In this work, we investigated interactions between the host and a gut pathogen across generations using the *Drosophila melanogaster* model. We found that pathogen density can influence intergenerational responses due to impacts on both the microbiome and on its ability to form low-level chronic infections within the host. Interestingly, these interactions have important impacts on non-immune physiological responses.

## Results

### Parental infection leads to pathogen persistence without altering microbiome composition

To explore the effects of a gut pathogen on host immunity and the microbiome across generations, we infected a parental generation of *Drosophila* females and males using three different doses of *Pseudomonas entomophila* (*Pe*), a high (OD100), medium (OD50), and low (OD5) dose. We assessed survival for seven days post-infection and measured microbiome and pathogen load at different time points post-infection (Fig 1A). Fly survival (Fig 1B) and pathogen load were dose-dependent in females and males (S1 Fig). Although pathogen load decreased over time, we observed *Pe* persistence in all doses (Fig 1C). We observed an initial increase in total microbiome abundance (4 h and 24 h) in flies ingesting a medium dose of *Pe*. However, total microbiome abundance was the same in all infected conditions when compared to uninfected controls by seven days post-infection (Fig 1D). Moreover, there was no impact of *Pe* infection on the ratio between the two main groups of bacteria that comprise the *Drosophila* microbiome, Lactic Acid Bacteria (LAB) and Acetic Acid Bacteria (AAB) (Fig 1E), suggesting infection did not alter microbiome composition. Overall, our result in the parental population indicates that, in terms of survival, females and males exhibit a dose-dependent response over time and even though there is a reduction in the pathogen load, *Pe* persists in the surviving parents. Despite shifts in the microbiome at early time points, the microbiome of the infected parental generation recovered in both abundance and composition, like that observed in uninfected controls.

**Fig 1.**
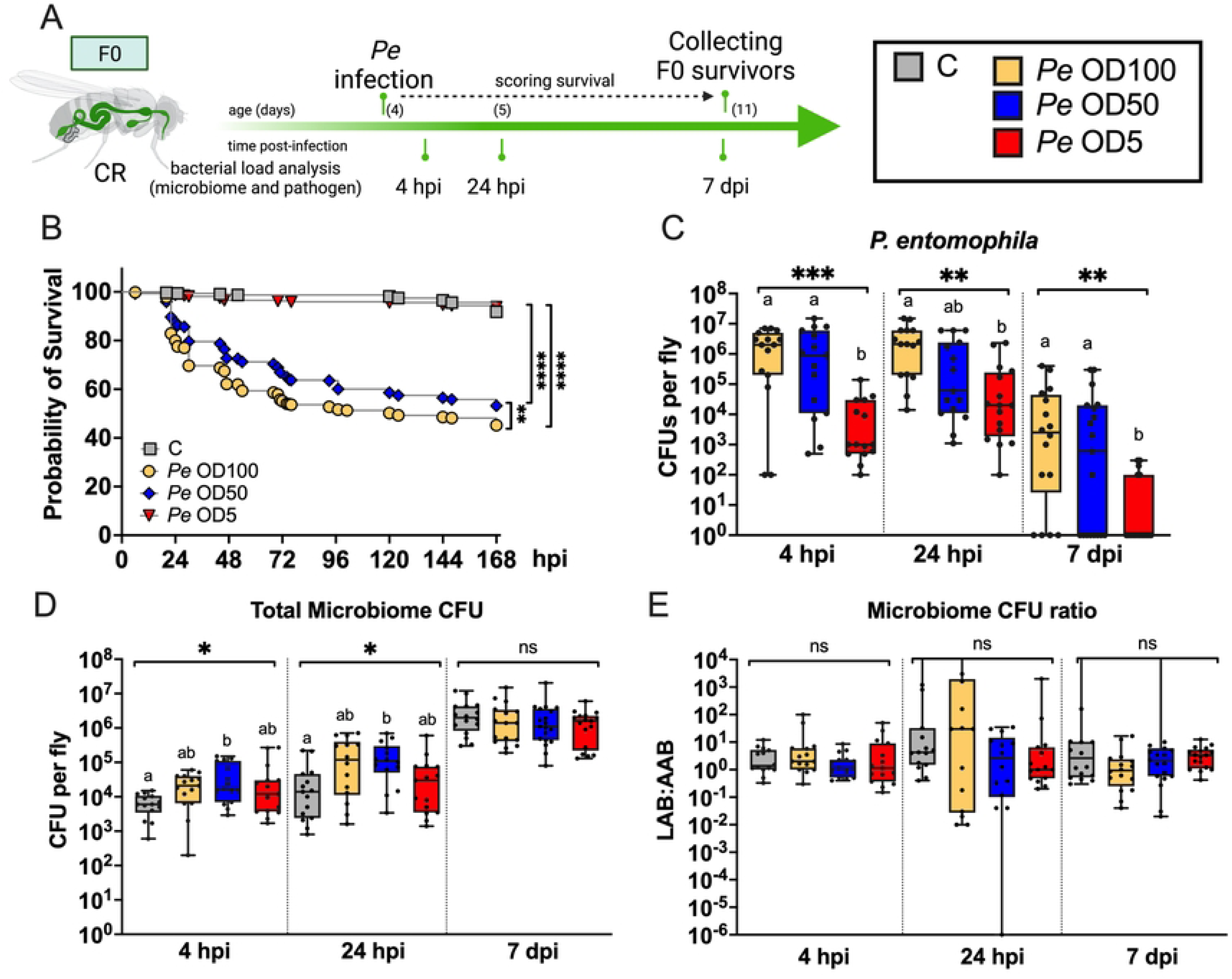
Pathogen persistence in the parental generation did not alter microbiome composition. **A)** Experimental strategy for the parental infection using *Pseudomonas entomophila* (*Pe*). **B)** Survival curve shows dose response to *Pe* infection. **C)** Bacterial load of the pathogen showed dose effect and persistence of the pathogen in the survivors at 7 dpi. **D)** *Pe* infection increased total microbiome at early time points post-infection, but microbiome is re-established at seven dpi. **E)** The microbiome ratio between Lactic Acid Bacteria (LAB) and Acetic Acid Bacteria (AAB) was not altered after *Pe* infection. Statistical analyses were performed using data over three biological replicates. Log-rank (Mantel-Cox) test for survival (B), and Kruskal-Wallis followed of Dunn’s multiple comparisons test were used for bacterial load analysis (C-E). Box plots indicate the median, and black points represent each data point; whiskers go to the minimum or maximum. Significance is expressed as follows: ns, not significant (p > 0.05); ****p =< 0.0001; ***p < 0.001; **p < 0.005 and, *p < 0.05. Values labeled with different letters are significantly different in Dunn’s multiple comparisons test. Details about the number of flies used and statistical values are found in S1 Table.

### Parental infection leads to environmental transmission of a pathogen to offspring

In nature, as in lab conditions, “*Drosophila* lives where it eats,” oviposition, hatching, and larval development occur on the same substrate (25). This shared living environment means that pathogen experiences of the parents have the potential to impact future generations directly through exposure to the same pathogens. Given the persistence of *Pe* in the parental generation (Fig 1C), we assessed whether the pathogen could also persist in subsequent generations. To test this, we collected all surviving adults seven days post-infection and used these to establish populations for each dose, as well as a control population, and followed the offspring for two generations. We sampled the food substrate at the start (S) and end (E) of F1 and F2 development (larval rearing environments) and sampled flies four days post-eclosion to quantify bacterial load (Fig 2A). Parents, regardless of infection dose, transferred *Pe* to the food in which the F1 were developing, and *Pe* load in food increased across development time (Fig 2B). This growth in the substrate resulted in F1 offspring of parents challenged with the high (OD100) and medium (OD50) *Pe* doses being exposed to similar pathogen loads during development. In contrast, F1 offspring from parents challenged with *Pe* OD5 (low dose) developed in an environment with a lower pathogen load (Fig 2C-D). However, the level of pathogen prevalence and load offspring experienced as larvae did not necessarily dictate adult conditions. As adults, F1 offspring exposed to the high *Pe* dose exhibited the highest *Pe* prevalence and pathogen load. Whereas, as adults *Pe* OD5 F1 offspring exhibit low pathogen prevalence and load comparable to *Pe* OD50 F1 offspring, indicating that persistence in offspring does not depend on the capacity of the pathogen to grow in the food environment.

**Fig 2.**
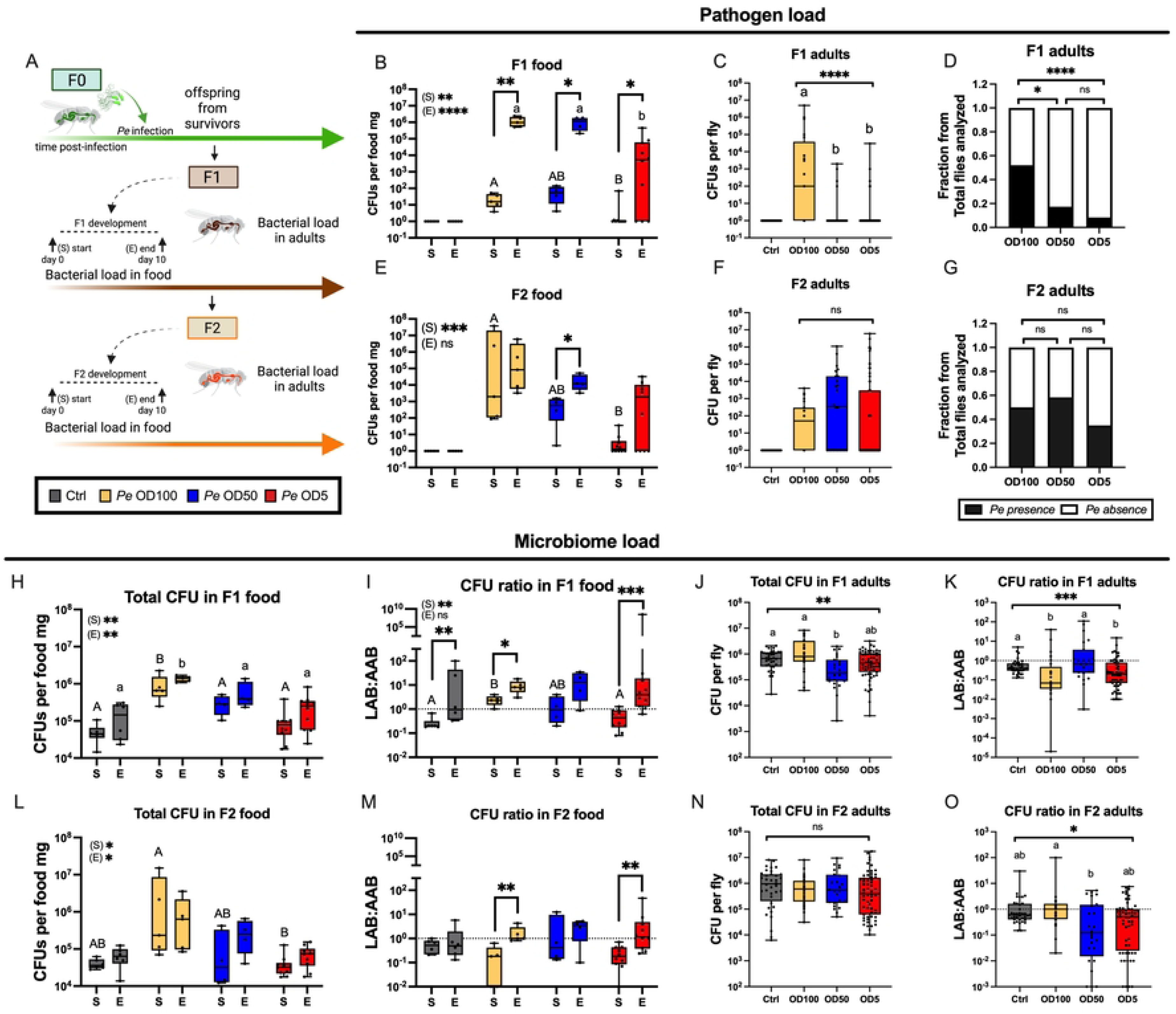
Environmental transmission of a pathogen led to microbiome remodeling across generations. **A)** Experimental strategy to study parental infection effects across generations during *Pseudomonas entomophila* (*Pe*) infection using three different doses: *Pe* OD100 (High), *Pe* OD50 (Medium), *Pe* OD5 (Low). Samples were taken from the food at the start (S) and end (E) of the F1 and F2 development. Four-day-old flies were used to quantify the bacterial load. **B)** Pathogen load in the food across the F1 development. **C)** Pathogen load and **D)** proportion of infected flies in F1 adults. **E)** Pathogen load in the food across the F2 development. **F)** Pathogen load and **G)** proportion of flies infected in F2 adults. **H)** Microbiome alterations in F1 food and **L)** F2 food. **I)** Microbiome remodeling between Lactic Acid Bacteria (LAB) and Acetic Acid Bacteria (AAB) in F1 and **M)** F2 food. **J)** Alterations of total CFU and **K)** microbiome CFU ratio in F1 adults. **N)** Total microbiome was not altered in F2 adults. **O)** Microbiome remodeling in F2 adults. Box plots for graphs showing bacterial load data indicate median and black points representing one fly at each point; whiskers go to the minimum or maximum. Significance is expressed as follows: ns, not significant (p > 0.05); ****p =< 0.0001; ***p < 0.001; **p < 0.005 and, *p < 0.05. Mann Whitney test was used to compare values between the S (start) and E (end) of the experiment and significant values are shown on the graph. Kruskal-Wallis test followed by Dunn’s multiple comparisons test was conducted to compare values according to the parental treatment in each generation per time. Significant values across the time scale are indicated next to (S) or (E). Treatments labeled with different letters are significantly different in Dunn’s multiple comparisons test, with upper-case letters for (S) and lower-case letters for (E). A detailed version of the experimental strategy, LAB, and AAB per treatment data is indicated in S2 Fig. Three biological experiments were conducted using 1-3 vials depending on parental survivors, details about the number of flies used and statistical values are found in S1 Table.

The transmission and persistence of *Pe* was also observed in the second generation (F2) as offspring developed (Fig 2E). Pathogen load in the F2 food at the start of the experiment was dose-dependent (Fig 2E). Yet, by the end of the development period *Pe* loads reached similar levels in the food across all treatments, such that F2 offspring were exposed to similar pathogen loads during development (start vs. end), which led to similar pathogen load and prevalence at the adult stage (Fig 2F-G). Together, our data show that the persistence of a pathogen in the parents led to environmental transmission to offspring across subsequent generations, and the dose of the pathogen to which the parents were exposed impacted the offspring environment. These findings also indicate that prevalence and persistence in offspring may be independent of the ability of the pathogen to grow in the food environment.

### Environmental transmission of a pathogen alters offspring microbiome composition

The microbiome significantly influences general host physiology, including development and immunity. Given our observation that the parents transmitted the *Pe* pathogen to the offspring through food, we next explored whether pathogen exposure impacted the microbiome in subsequent generations. Total microbiome load of F1 *Pe* OD100 offspring increased compared to F1 offspring from uninfected parents or offspring of F1 medium or low treatments. In contrast to pathogen load, total microbiome counts were stable in F1 food across development for all treatments (Fig 2H). Although total microbiome counts were stable, the microbiome ratio between lactic acid bacteria (LAB) and acetic acid bacteria (AAB) shifted over development time in F1 food regardless of parental treatment (Fig 2I).

Microbiome counts in F2 food followed similar trends to those observed in F1 food, though total microbiome loads were similar across conditions and developmental time (Fig 2L). The microbiome ratio between LAB and AAB increased in the food, but only where F2 *Pe* OD100 and *Pe* OD5 offspring developed (Fig 2M). In contrast to food, microbiome alterations in F1 and F2 adults were specific to parental treatment. For example, F1 adults from *Pe* OD50 treated parents had a lower microbiome load than uninfected flies, but in F2 offspring, the total microbiome is similar between groups (Fig 2J, N). Moreover, LAB abundance was altered in both food and adult offspring, causing a shift in the ratio between LAB and AAB in F1 and F2 adults (Fig 2K, O). However, the LAB abundance decreases in the offspring from *Pe* OD5 parents in both generations (S3 Fig). These findings indicate that parental infection affects the microbiome status in subsequent generations. Our results further suggest that the persistence of *Pe* in the food and the offspring significantly contributes to such microbiome alterations.

### Pathogen persistence correlates with pathogen virulence

Considering that the pathogen persists in the food where the offspring develops, and such offspring are chronically exposed to the pathogen, we next assessed the virulence of *Pe* isolated from the food and adult flies. The GacS/GacA two-component system globally regulates nearly all of the currently identified *Pe* virulence factors (i.e. pore-forming toxins, proteases, etc.), including the metalloprotease *AprA* (26). As such, GacS/GacA mutants can be identified by their loss of protease activity when cultured on skim milk plates. We detected positive protease activity in all the *Pe* isolated from F1 food (Fig 3A). Protease active isolates were also isolated from *Pe* OD50 and *Pe* OD5 F1 offspring as adults (Fig 3B). However, no protease active isolates were cultured from *Pe* OD100 F1 flies. Interestingly, the pathogen load and the proportion of total flies with detectable *Pe* are higher in *Pe* OD100 F1 flies (Fig 2C-D), suggesting that the loss of virulence aided persistence in the adult gut across generations.

**Fig 3.**
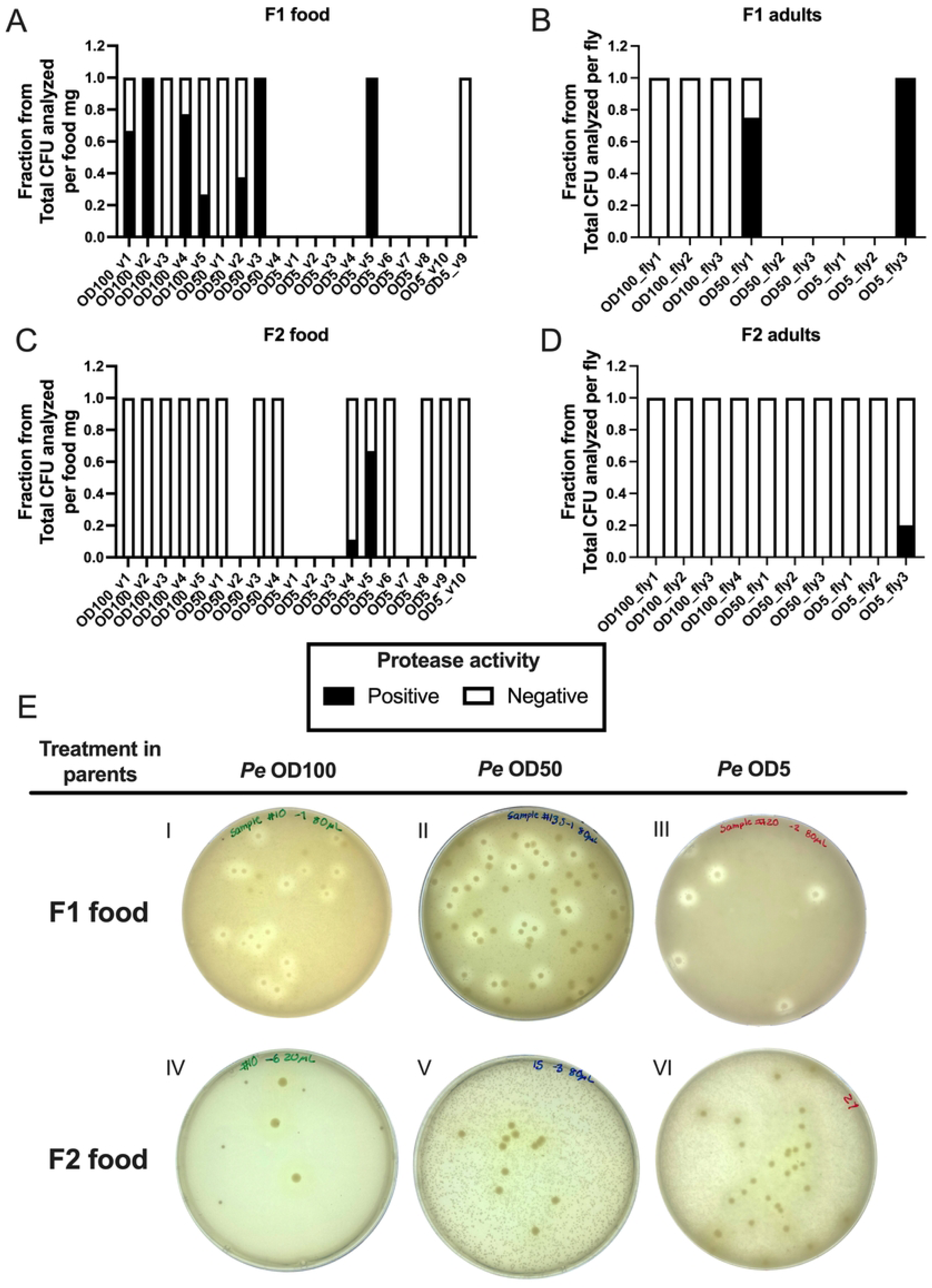
Protease activity of *Pseudomonas entomophila* in the food and the offspring differs across generations. The virulence of *Pe* found in food and adult offspring samples from each generation was explored by testing protease activity on skim milk plates. **A)** Protease activity was present in the food where F1 developed. **B)** In F1 adults, bacteria isolate from *Pe* OD100 offspring showed absence of protease activity. **C)** Protease activity was absent in most of the F2 vials analyzed during the study. **D)** In F2 adults, protease activity was detected only in bacteria isolated from *Pe* OD5 offspring. **E)** Representative images showing visible protease activity of *Pe* on skim milk plates from F1 food (I-III) and F2 food (IV-VI). The absence of a bar in the graphs indicates that *Pe* was not detected in the sample.

In contrast to F1, protease activity is absent in *Pe* isolates from food in which *Pe* OD100 and *Pe* OD50 F2 offspring develop (Fig 3C), we only detected protease activity in bacteria isolated from the food from *Pe* OD5 offspring. Similarly, *Pe* isolated from F2 adults lacked protease activity, and only *Pe* from *Pe* OD5 offspring retained protease activity (Fig 3D). These findings support that, in addition to the differences found in the pathogen load and prevalence in flies infected in F1 and F2, the loss of pathogen virulence is correlated with greater persistence across generations.

### Chronic pathogen exposure has varied impacts on host physiology across generations

Hosts can regulate the activation or inhibition of immune responses to match infection intensity to conserve energy. However, even when the intensity of the infection is very high, other fitness components may be prioritized, leading to a reallocation of energetic resources, for example by investing in reproduction instead of survival (27). In addition, chronic infections are energetically costly (28), as carrying the pathogen for longer time periods may compromise survival to additional stressors, such as food limitation. We hypothesized that long-lasting exposure to the pathogen in the parents and then offspring would lead to several fitness consequences. To assess this, we explored pupal development success, fecundity, and the starvation resistance response in both parents and offspring across generations (Fig 4A; see S3A Fig for details). Infection altered parental fecundity in a dose-dependent manner, which impacted total pupae numbers of the F1 generation (Fig 4B), but not pupal eclosion rates (Fig 4D), indicating that if an offspring reached the pupal stage, it had similar eclosion success regardless of parental infection status. Notably, in the second generation (F2), the effect of parental treatment was inverse to that of the F1 (Fig 4C). Total pupae per fly was highest in the offspring from *Pe* OD100 and *Pe* OD50 infected parents compared to *Pe* OD5 and uninfected parents. Interestingly, pupae of *Pe* OD100 infected parents exhibited higher eclosion rates (Fig 4E). The higher eclosion rates correlated with the presence of less virulent pathogens in the second generation, suggesting that *Pe* may disrupt food nutrient acquisition and impact larvae development and pupae eclosion. Such results also indicate that parental infection exerts reproductive and developmental impacts that extend at least two generations.

**Fig 4.**
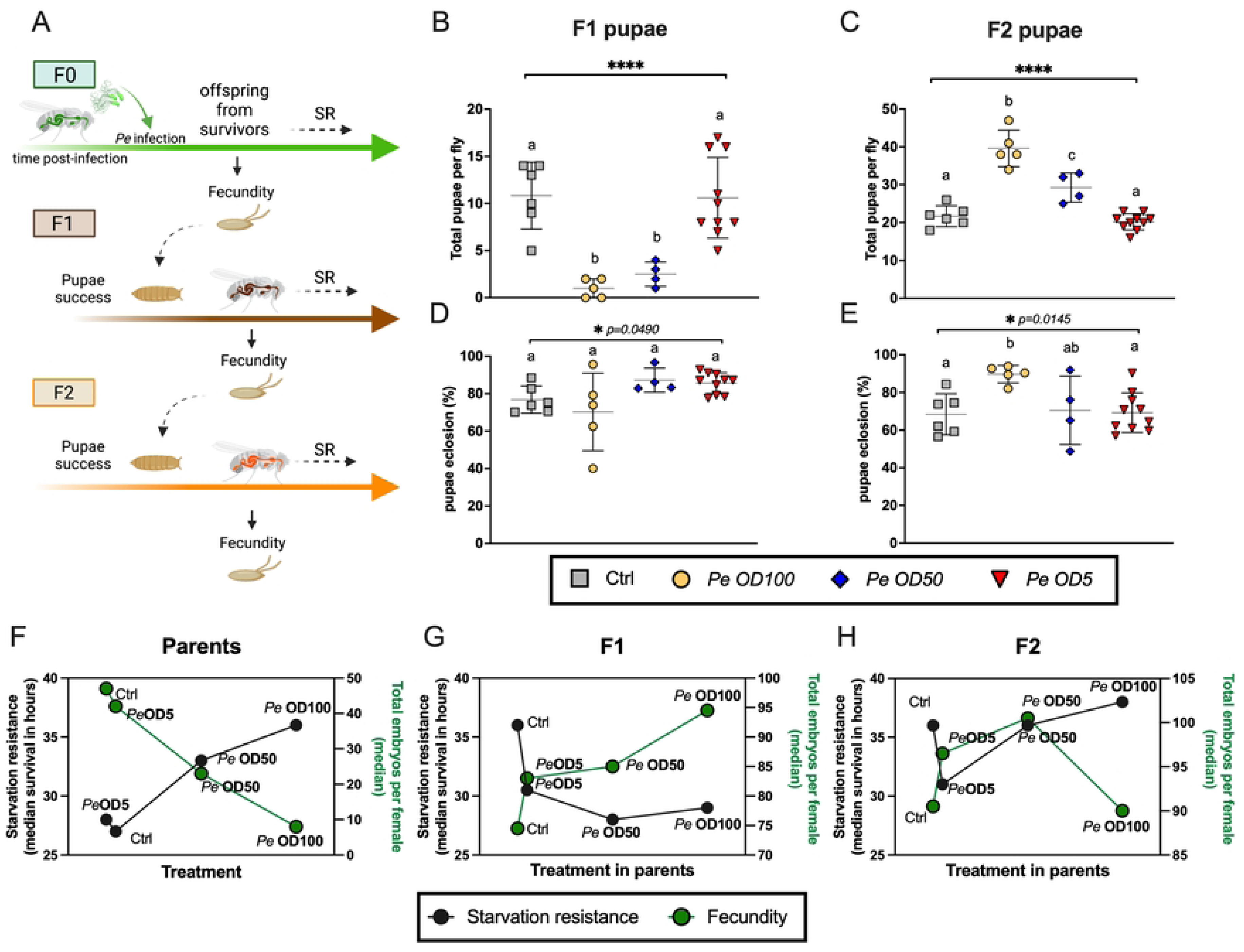
Chronic exposure to the pathogen in the parents and the offspring led to fitness consequences. **A)** Experimental strategy to study fitness consequences during chronic pathogen exposure. Pupae development success and fecundity was scored in each generation. In addition, starvation resistance response was evaluated in the parents and offspring. For details on the experimental strategy, see S3A Fig. **B)** Total pupae per fly in F1 offspring. **C)** Total pupae in F2 offspring showed a positive effect according to the infection dose used in the parental treatment. **D)** Parental infection had a marginal effect on pupae eclosion in F1 pupae. **E)** In F2, offspring from *Pe* OD100 parents showed increased pupae eclosion. **F)** Fecundity and sensitivity to starvation in the parents (F0) showed an inverse relationship with *Pe* infection in a dose-dependent manner. **G)** Fecundity and sensitivity to starvation resistance in F1 **H)** and F2 offspring. In graphs B-E), each point represents data from one vial. Graph shows mean and standard deviation. One-way ANOVA followed Tukey’s multiple comparisons test was used to determine significant differences between groups. Significance is expressed as follows: ns, not significant (p > 0.05); ****p =< 0.0001; ***p < 0.001; **p < 0.005 and, *p < 0.05. Pearson correlation for linear association between fecundity and starvation resistance response are indicated in S4 Fig. Graphs show data from three biological replicates with 1-3 vials depending on parental survivors’ availability, details about the number of flies used and specific statistical test applied to compare fecundity and starvation resistance between groups are found in S1 Table.

In examining starvation-resistance of parents and offspring, we observed that infection resulted in parents being less sensitive to starvation in a dose-dependent manner (Fig 4F, and S3B-C Fig). The increased survival in response to starvation observed post-infection inversely correlated with fecundity (S4A Fig; Pearson correlation, r= −0.9677; p=0.0323), suggesting a trade-off between fecundity and starvation resistance. These results also suggest that energetic resources were reallocated in the parents between these physiological functions during *Pe* infection. In F1 offspring, the effects were less clear. F1 offspring exhibit roughly a dose-dependent impact on fecundity, but there was no dose effect on their starvation resistance and instead offspring from parents infected with the high and medium dose were more sensitive to starvation than uninfected offspring (S3E and S4B Fig; Pearson correlation, r= −0.8065; p=0.1935). Overall, in F2 the dose-response impact on starvation resistance was like that of parents. However, the fecundity did not correlate with this response (S4C Fig; Pearson correlation, r= −0.4292; p=0.5708). We only observed a trade-off effect in the offspring from *Pe* OD50 parents, which was more sensitive to starvation but had higher fecundity (S3F-G Fig) when compared to offspring from uninfected parents. Altogether, our results show that the infection caused physiological changes in parents and offspring, and those changes may depend on the dose, suggesting a cost to pathogen tolerance across generations.

### Prior *Pe* infection does not confer protection to subsequent infection

Some insects transfer protection to their offspring when infected with a pathogen. To begin evaluating the effects of the parental infection on offspring, we challenged adult F1 and F2 offspring from all parental treatments with a high dose (OD100) of *Pe* to assess survival (Fig 5A and S5 Fig for detail). There was no significant difference in offspring survival after a new *Pe* challenge in either generation (Fig 5B-C). Due to the transmission of *Pe,* we detected pathogen in the unchallenged offspring treatments (control groups fed with sucrose solution, but carrying the pathogen because of parental infection). However, the density of *Pe* in these treatments was quite variable across time (S6 Fig). For the offspring treatments newly challenged with *Pe*, the pathogen load of F1 and F2 offspring was the same across treatments independent of parental challenge (Fig 5D-G; S6 Fig). These data indicate that chronic exposure to *Pe* did not provide an advantage upon a new challenge with the same pathogen.

**Fig 5.**
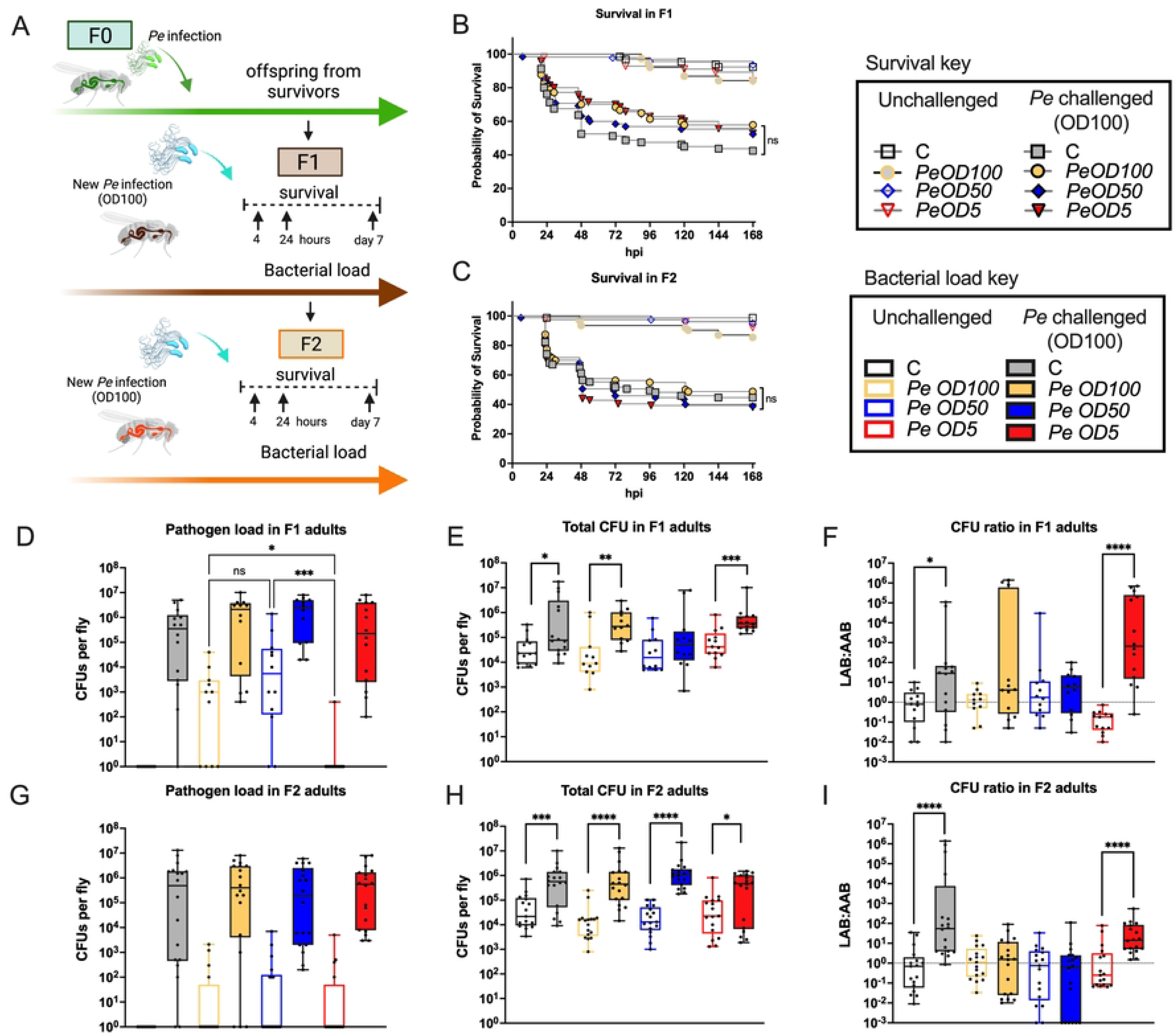
Chronic exposure to the pathogen in the offspring did not improve survival but did cause microbiome remodeling after a new infection. **A)** Experimental strategy to explore offspring response to a new infection. Parents were infected using three different doses, *Pe* OD100 (High), *Pe* OD50 (Medium), and *Pe* OD5 (Low); offspring were collected and reared to reach the adult stage. Adults from each generation were challenged (*Pe* challenged) using a high dose of *Pe* (OD100). Survival and bacterial load were determined for each generation. For experimental strategy details see S5 Fig. **B-C)** Survival curve after the new *Pe* challenge showed similar survival independent of the treatment in the parents in both generations. Pathogen load in F1 offspring **(D)** and F2 offspring **(G)**, after the new challenge with *Pe.* E) Offspring from parents infected with the medium dose did not show changes in microbiome abundance in F1 but showed altered microbiome in F2. **H)** Abundance of microbiome members increased independently of the parental treatment when newly challenged with *Pe.* F, I) CFU ratio between Lactic Acid Bacteria (LAB) and Acetic Acid Bacteria (AAB). Graphs show bacterial load data at 24 hpi; box plots indicate median and black points represent one fly per data point, and whiskers go to the minimum or maximum. Statistical analyses were performed using the Log-rank (Mantel-Cox) test for survival. Mann Whitney test was used to compare bacterial load between pairs unchallenged vs new *Pe* challenged in each parental treatment. For pathogen load, Kruskal-Wallis test, followed by Dunn’s multiple comparisons test, was conducted to compare values between parental treatment in each offspring treatment (between unchallenged only or between new *Pe* challenged only). Significance is expressed as follows: ns, not significant (p > 0.05); ****p =< 0.0001; ***p < 0.001; **p < 0.005 and, *p < 0.05. Data at four hours and seven days post-infection are indicated in S6 Fig. Specific statistical test and the number of flies used over three biological replicates are in S1 Table.

### Parental infection influences microbiome shifts in response to a new infection

One aspect that has received less attention in the study of immune response across generations is the impact of or on the microbiome. Yet, the microbiome impacts host immune status, which in turn modulates the microbiome. Chronic pathogen exposure altered the microbiome, particularly in terms of composition (ratio between LAB and AAB (Fig 2)). We next evaluated the impact of a new *Pe* challenge on the microbiome. In both F1 and F2 offspring, we observed that a new *Pe* infection increased microbiome load, though this was not significant for F1 offspring from parents infected with the medium dose (*Pe* OD50) (Fig 5E and 5H). This increase in microbiome abundance lasted up to seven dpi in offspring from parents infected with *Pe* OD5 (S6 Fig), suggesting the effects of infection can be long-lasting under some conditions. In terms of composition, new *Pe* challenge increased the number of LAB in offspring from uninfected parents and offspring from parents infected with the low dose (*Pe* OD5), but there was no significant shift in composition in offspring from *Pe* OD100 and *Pe* OD50 parents (Fig 5F, and I). Our results indicate that a new *Pe* infection causes a shift in the ratio between LAB and AAB in F1 and F2 adults, and such effects are influenced by the infection experience of the parents.

### Chronic pathogen exposure alters gene expression of F1 and F2 offspring

In *Drosophila,* gut bacterial infections trigger an immune response, upregulating the expression of antimicrobial peptides (AMPs). Dysregulation of the immune response can cause physiological changes in the host, altering homeostasis. We explored the possibility that parental exposure and chronic exposure to the pathogen may lead to changes in offspring immune responses. We quantified the expression of genes associated with host responses to *Pe*, including the expression of AMPs, such as Diptericin A (*DptA*), Attacin A (*attA*), Metchnikowin (*Mtk*), Drosomycin-like 3 (*Dro3*); stress response molecules including Dual oxidase (*Duox*), Glutathione S transferase D5 (*GstD5*), and Growth arrest and DNA damage-inducible 45 (*Gadd45*); and signaling molecules involved in maintaining proper gut homeostasis, for example JAK/STAT pathway components as unpaired 3 (*upd3*), Suppressor of cytokine signaling at 36E (*Socs36E*), and JNK signaling such as puckered (*puc*). We found that new *Pe*-challenge in F1 offspring of *Pe*-challenged parents (independent of dose) exhibited increased expression of AMPs, stress response molecules, and JAK/STAT, JNK pathway components compared to control *Pe*-challenged flies at the early time of the infection (4h) (Fig 6A). Interestingly, F1 unchallenged offspring from *Pe*-challenged parents have suppression in the expression of most of the genes evaluated, except for *Gstd5*. The upregulation of *Gstd5* correlates with the presence of protease active pathogen in F1 offspring, suggesting that detoxification mechanisms are activated to reach a homeostatic balance of the host in the first generation.

**Fig 6.**
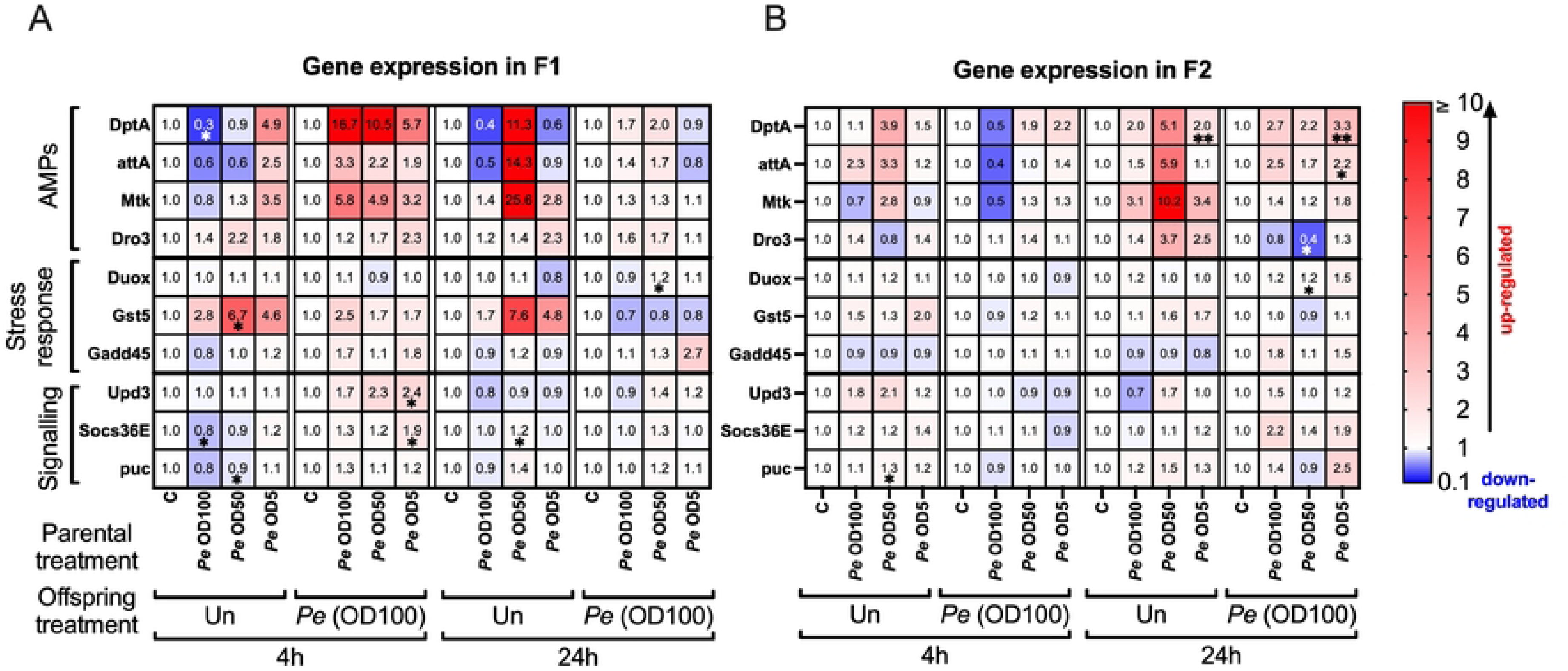
Differential gene expression between F1 and F2 offspring during early response to the new *Pe* infection. Gene expression in F1 **(A)** and F2 **(B)** offspring after the new *Pe* infection. Heat map shows the mean of the mRNA fold change in the unchallenged and newly *Pe* challenged at 4 and 24 hpi. Expression was normalized to the control treatment (unchallenged condition in the parents) for each time and each offspring treatment (value 1, shown in white). Up-regulated values are shown in red, and down-regulated values are shown in blue. Numbers indicate mean values from three biological experiments. An unpaired T-test with Welch correction was used to determine significant differences between the control treatment in the parents and the dose used in each parental treatment following a new *Pe* infection in offspring. Significance is expressed as follows: **p < 0.005 and *p < 0.05. For details in graphs for each gene per treatment, see S6 Fig.

In F2 offspring, *DptA* and *attA* expression were significantly up-regulated only in the offspring from *Pe* OD5 parents at 24 hpi (Fig 6B). The increased expression observed in the F2 *Pe* OD5 offspring is significant compared to the control *Pe*-challenged offspring. However, such expression does not improve survival or pathogen clearance. In contrast with the offspring from the other parental treatments, the increase in *DptA* and *attA* expression is correlated with the presence of protease active pathogen found during the adult stage in F2 *Pe* OD5 offspring, suggesting that the virulence of the pathogen may affect the capacity to induce AMPs gene expression in future pathogen encounters. Although the increased immune response is not beneficial for the host in terms of survival during the new immune challenge, it might have selective immunomodulatory effects derived from the parental infection in non-immunity-related function during host-pathogen adaptation.

## Discussion

*Drosophila* has been a valuable model for studying host-microbe interactions in both beneficial and pathogenic contexts, and the effects of such interactions impact several aspects of host physiology, including immunity, reproduction, and metabolism. However, most studies have concentrated on such impacts within a single generation. Here, using the *D. melanogaster* model, we studied the impact of parental infection on host physiology and immunity across generations. Our study shows parental *Pe* infection led to the persistence of the pathogen and its transmission to subsequent generations, in a dose-dependent manner. Yet, pathogen virulence in subsequent generations was inverse of dose. Such pathogen virulence and density can influence host physiological responses in the offspring due to impacts on the microbiome and on its ability to form low-level chronic infections.

### Pathogen virulence as a crucial factor for persistence

Our study showed that pathogen load changed across generations (Fig 7A) in a manner dependent on initial parental infection dose. However, we observed that pathogen virulence changed in the food environment and in the fly, wherein an initial high dose led to a less virulent pathogen in the next generation (Fig 7B). *Pe* virulence depends on genes controlled by the GacS/GacA kinase system. This signal transduction system regulates the expression of virulence factors, including extracellular enzymes, such as the metalloprotease *AprA*. Mutations in GacA are avirulent, reported to persist less in flies, and fail to activate the immune response (29). Whereas, *Pe* with a mutation in the gene encoding the metalloprotease *AprA (Pe AprA* mutants) are reported to show attenuated virulence, persist better than *Pe gacA* mutants, and induce host immunity (26). In F1 and F2 generations, we observed the presence of protease-negative isolates, particularly from flies fed the high and medium *Pe* doses. This indicates that despite our observation of chronic carriage of the pathogen in all doses, the high and medium dose offspring were exposed to a higher density of less/non virulent pathogens. As such, *Pe* virulence shapes its ability to persist in the fly and the environment (Fig 7C). Our study also showed that chronic exposure of offspring to *Pe* altered their immune response in a manner dependent on virulence and density (Fig 7D). For instance, offspring from parents challenged with a high dose, suppressed their immune response in the first generation, evidenced by the downregulation of genes generally induced by *Pe*. In contrast, the offspring from parents challenged with medium and low doses, in which the *Pe* prevalence was lower, but the proportion of protease-positive, meaning virulent, *Pe* was higher, the immune response was upregulated. These results suggest that virulence factors are probably related to the regulation of the immune response observed and it also likely reflects impaired gut epithelium renewal and immunity that contribute to the persistence of the pathogen in subsequent generations. Considering our results, we hypothesize that across generations, there is a balance between host factors and pathogen factors for mutual survival of the host and the pathogen; the host immune response is shaped by pathogen virulence, where low virulence prevents too strong of an immune response, allowing pathogen survival. This balance suggests a tolerance mechanism that allows both pathogen persistence in the environment and the success of subsequent host generations.

**Fig 7.**
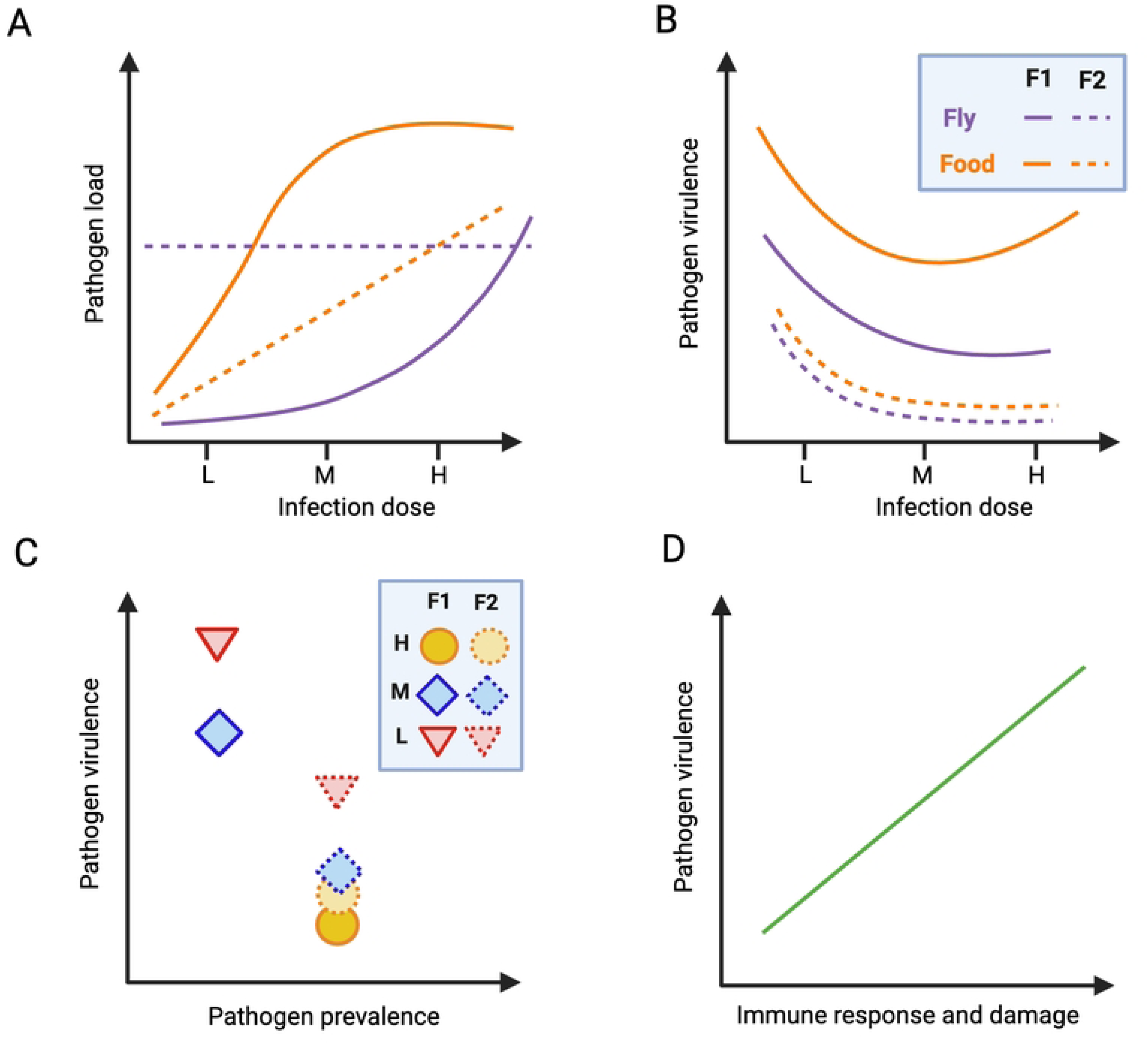
Summary of the effects on the pathogen and host immunity across generations. **A)** The infection dose impacted the pathogen load in both food and flies. **B)** Differences in pathogen virulence over generations depend on the initial pathogen dose. In F2, high- and medium-dose offspring are exposed to less virulent pathogens. **C)** Pathogen virulence correlates with pathogen prevalence in the fly. **D)** Chronic pathogen exposure alters the host immune response depending on pathogen virulence and density.

### Insights of overlapping networks between host physiology and survival

One important question raised by our study is whether chronic pathogen exposure in offspring impacted other aspects of host physiology, such as reproduction and survival under stress conditions. Infection and starvation are commonly driven by similar networks and resources. Both modulate metabolism with the aim of maximizing the likelihood of survival (30). Moreover, reproductive activity and egg production in *Drosophila,* is generally constant across adulthood (with the exception of age-dependent effects), indicating that the processes of egg maturation and laying require a constant demand and expenditure of energy (31,32). Our study showed a negative correlation between fecundity and starvation resistance indicating that *Pe* infection induces a dose-dependent trade-off in the parents. Interestingly, it has been shown that infection in parents can lead to prioritizing survival and investment in the immune response over egg production, suggesting a trade-off between reproduction and survival under stress conditions (33). In the same manner, our work supports that such trade-offs can influence subsequent generations. We also observed that the trade-off in offspring is different depending on pathogen dose and virulence. When we observed higher pathogen virulence, most host resources were put to the immune response; when virulence is low, most resources were shunted to reproduction, possibly to compensate for reduced survival when developing in the presence of the pathogen. In the second generation, when pathogen virulence was low, and prevalence was higher than the first generation, we observed shifts in reproduction and stress responses, suggesting resource usage as in uninfected conditions. The host physiological effects we observed are likely influenced by the route of infection used in our study. A previous study using *Drosophila* populations co-evolved with *P. entomophila* following systemic infection (direct injection into the body cavity) found no differences in fecundity or other life history traits and concluded that after multiple generations, the host and pathogen adaptation did not have a physiological cost (34). In contrast, our work found that following oral infection with *Pe* there were multiple physiological impacts on the host, within and across generations. This makes sense, given that enteric infection has the capacity to impact many aspects of host physiology, and not only host immunity. Moreover, we consider that physiological responses involving a reproductive or energetic cost, are a result of an adapted response to infection that may contribute to the tolerance of the population that balances pathogen persistence over generations and the host survival.

### Microbiome remodeling impacts host physiology across generations

Previous work in *Drosophila* has shown that the microbiome has a crucial role during infection with *Pe,* modulating the gut immune response to infection (11,35). Moreover, in other insect-microbe interactions, it has been shown that the composition of the bacterial community in the parents and transmission of such bacterial community to the offspring, influences several aspects of offspring performance, including larval growth and development, immune response and gene expression (36–38). Our findings indicated that microbiome remodeling induced by *Pe* can affect subsequent generations via alterations of the food environment, and such microbiome remodeling is relevant to the experience of offspring with the pathogen. We observed that LAB abundance decreased in offspring of parents challenged with a low dose in both generations; however, when challenged newly with a high dose of *Pe*, the offspring exhibited an increase in total microbiome load and LAB density. We suspect that this effect is related to the proposed role of the microbiome in colonization resistance, wherein the microbiome can act as a physical barrier in the gut to limit pathogens (12). Flies associated with *L. plantarum* have shown improved survival and limited gut colonization by a number of Gram-negative pathogens, including *Pe* (35,39). In our study, the increased bacterial density did not improve survival or bacterial clearance across the generational time scale we evaluated, highlighting that the context of association may be important for such effects. Future research is needed to understand if microbiome remodeling impacts other physiological aspects of the host, such as life span, or if microbiome function is changed. In addition, the microbiome might directly impact the pathogen, altering virulence. Understanding these microbe-microbe interactions and what factors might mediate such effects is of great interest.

Alternatively, it is also possible that pathogen virulence and persistence in the adult stage are related to beneficial or harmful aspects of host physiology observed during the offspring development by disrupting food nutrient acquisition and impacting pupae eclosion. In the second generation, during development, the *Pe* present was largely avirulent, as measured by the loss of protease activity. In such circumstances, it could be that the avirulent *Pe* is now functioning more directly as food or indirectly in making more nutrients available from the food, explaining the increased pupae eclosion we observed in the offspring. Previous studies have shown that microbiome members contribute nutrients to the host, thus modulating nutrient physiology (40,41). Future research is needed to understand whether virulent or avirulent pathogens have differing impacts of nutrient resources in the food substrate, where multiple generations interact with other microbes and pathogens. Overall, our observations highlight that during chronic pathogen exposure, the source and route of infection, dosage, virulence state, and microbiome composition are all relevant to understand the physiological consequences in the host.

### Beyond host protection during subsequent infections

In insects, previous interactions of larvae with a pathogen have been shown to influence adult susceptibility. For example, in mosquitoes, adults of previously infected larvae exhibited increased immune responses, although this did not increase their survival to infection as adults (42). In our experiments, offspring in both generations were chronically exposed to the pathogen, albeit with different levels of virulence, and yet we did not observe an improvement in survival or evidence of any protective effect to challenge. Thus, prior pathogen interactions are not necessarily beneficial or protective.

Our data indicate that prior parental pathogen exposure can alter the fitness of subsequent generations, though not necessarily through an immune-dependent mechanism. Offspring that developed in an environment of high pathogen density exhibited a decrease in the total number of individuals that developed to pupae, indicating a fitness cost in the first generation. However, in the second generation, pupae and eclosion success were higher than in the other doses or the control, indicating that larval development in an environment shaped by parental experiences can also affect the fitness of future generations. These host physiology and pathogen adaptations associated with the chronic interaction with the pathogen in the environment may be beneficial for the host in the long term. Infection increases the chance of pathogen spread throughout their niche, which may then infect their offspring and others in the population. We did not observe protection across generations to subsequent *Pe* infection, but instead found that *Pe’s* virulence was reduced across generations, allowing the insect to be chronically associated with an avirulent form of the pathogen. This outcome was unexpected and interesting, as it suggests that some aspect of persistent association with the host selects for loss of virulence. We do not yet know what the selection pressure was in our system, though we did see an inverse correlation between the initial dose of virulent pathogen and the proportion of avirulent pathogen in the next generation, suggesting that the level of host death in the population may be a contributing factor. It has been proposed that reduced virulence of a pathogen to a population will be selected for overtime given trade-offs between virulence and transmission (43). As seen in our study, it is possible that in nature, offspring of social insects or insects with gregarious behaviors may be more likely to be exposed to less virulent pathogens than when acquired from parents because the pathogens have already undergone modification due to host immunity. Thus, pathogen experiences might depend on pathogen state that in turn impacts several aspects of offspring physiology beyond the immune response.

Our findings are in agreement with previous work in *Drosophila* and other insects, that immune priming, or enhancement of the immune response, is greatly variable between insects (44–47). Such responses involve plasticity, depending on the pathogen, the infection route, and host status (48,49). In addition to the current criteria proposed to study TGIP (9,10), we propose that TGIP experiments should also consider changes in the pathogens, due to pathogen evolution, which is shaped by interactions with the host and other microbes in the environment.

In conclusion, our study highlights the dynamics of the host-pathogen interaction across generations. We did not observe evidence of immune priming or innate immune memory, as has been previously reported in some insect-microbe systems. Instead, in this work, we observed an interesting phenomenon of adaptation of the pathogen across generations, in which an initial high dose of virulent pathogen shifted to the loss of virulence, as evidenced by the loss of protease-positive isolates. Both mechanisms, immune priming, or pathogen adaptation/avirulence selection (in this work), confer a greater probability of host survival. Infection intensity and the pathogen virulence altered several aspects of host physiology that exerted different costs on the host, as a consequence of the shift from virulent to non-virulent invader (Fig 8). This is highly likely to occur in natural systems in other insect-pathogen interactions, and our work establishes a useful model to study such a phenomenon.

**Fig 8.**
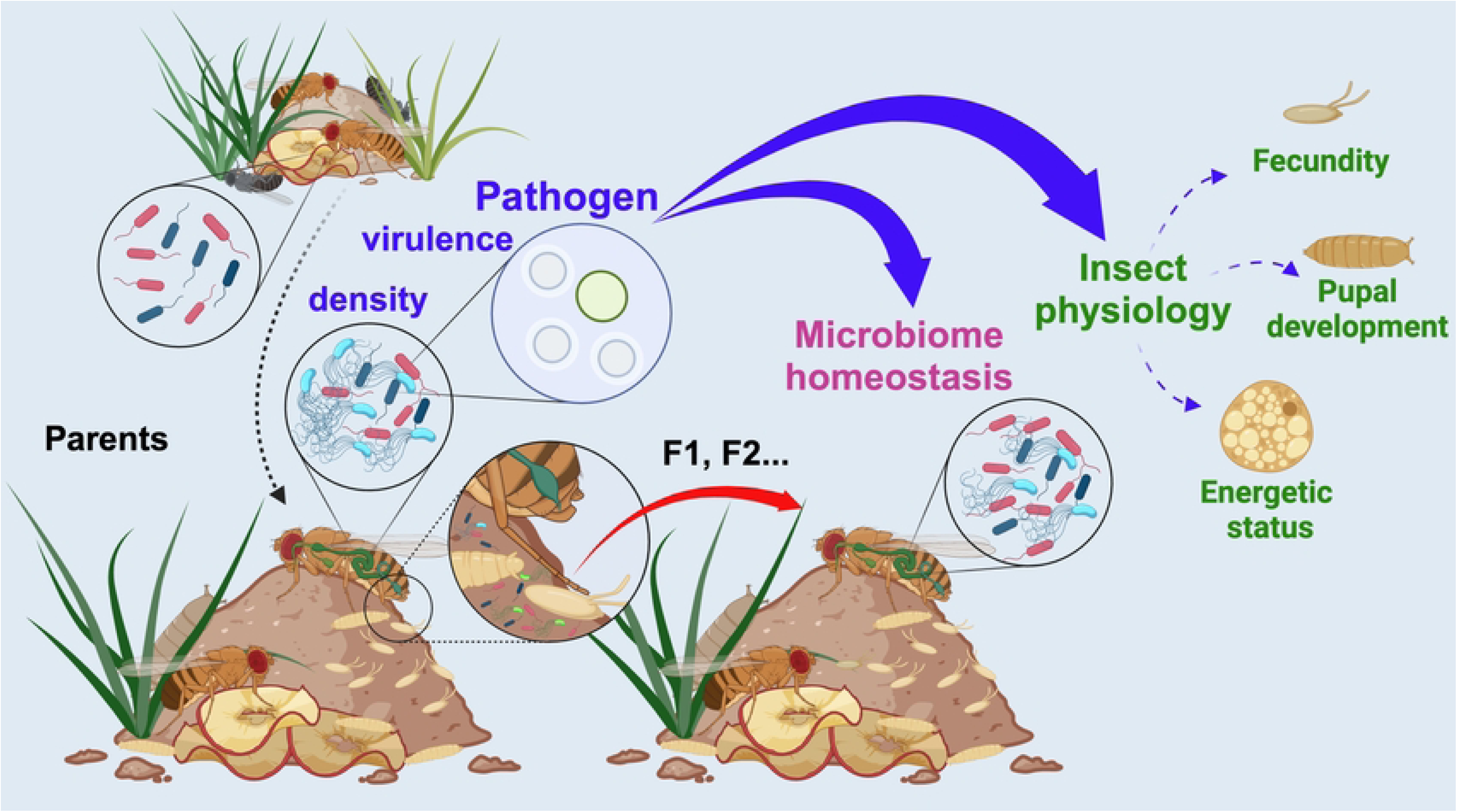
Model of the effects caused by *Pe* infection in the parents and in the subsequent generations. During *Pe* infection, the pathogen density and virulence are relevant for host response and physiology in the subsequent generations. Oral infection with *Pe* in the parents led to chronic exposure of the pathogen in the offspring with differences in virulence. Such chronic pathogen exposure altered the microbiome in offspring after the new challenge, depending on the dose of the parental treatment. Chronic exposure to the pathogen did not provide an advantage in survival during a new challenge and exerts a cost for several aspects of host physiology including reproduction, development, and energetic status as a consequence of the shift from virulent to non-virulent pathogen.

## Materials and Methods

### Fly stocks

CantonS flies (Canton Yale) were used for the experiments. *Wolbachia* was previously eliminated using tetracycline (0.05%). Fly stocks were maintained at a similar density (flies generated from 20 females and eight males per vial before starting any experimental procedure and during subsequent generations) on a 12:12 light: dark cycle at 60-80% relative humidity and 25°C. Flies were maintained on the Broderick Standard diet (components per liter of water: 40 g sucrose, 6 g agar, 50 g inactive dry yeast, 70 g yellow cornmeal, and 1.25 g methylparaben dissolved in 5 ml of 100% ethanol) (40).

### *Pseudomonas entomophila* culture method

For bacterial cultures, one *Pe* colony that was visually protease positive on Luria Bertani (LB) media (Invitrogen, #12795027) agar with 1% milk was used to start the bacterial culture on LB medium at 29°C for 18 hours. The bacterial pellet was collected by centrifugation, and protease activity was verified on LB-agar with 1% milk.

### Bacterial oral infection

Twenty-five flies were placed in empty vials for starvation for two hours at 29°C before being transferred to a food vial containing a paper disc. The bacterial solution was adjusted to a desired concentration by mixing equal volumes (1:1) of the bacterial pellet and a 5% sucrose solution (sterilized previously using a 0.22 µM membrane filter). The final concentration used per treatment was OD600 = 100 (high dose, *Pe*OD100); OD600 = 50 (medium dose, *Pe*OD50); and OD600 = 5 (low dose, *Pe*OD5) in 2.5 % sucrose, respectively. The bacterial solution was added to a desired concentration over the paper disc, and the flies were exposed to it for 24 hours; after that, the flies were transferred to a new sterile food vial. Flies were transferred to a new sterile food vial every two days and maintained at 29°C during the experiment. Mortality was monitored for seven days post-infection.

### Bacterial load analysis

A colony-forming unit (CFU) assay was performed to determine the bacterial load per fly. The fly body surface was sterilized by washing in 70% ethanol for 20 seconds, followed by washing with sterile PBS 1X, before transferring the fly to a 1.5 mL screw cap tube containing PBS 1X and 1.0 mm glass beads (BioSpec, #11079110). The sample was homogenized using the Bead Ruptor Elite homogenizer (Omni International, #1904E) at 4.0 m/s for 30 seconds. Serial dilutions of the homogenate were conducted manually using PBS 1X. The number of colonies on LB agar plates and DeMan, Rogosa, and Sharpe (MRS) agar media (RPI, #L11000) was determined after one and two days of incubation, respectively, at 29°C.

### Determination of bacteria in fly food

A sample of 10-15 mg of food was taken from the top of the vials and collected in a tube containing PBS 1X and 1.0 mm glass beads. The sample was homogenized twice using the Bead Ruptor Elite homogenizer at 4.0 m/s for 30 seconds. Serial dilutions of the homogenate were conducted manually, and the number of colonies on LB agar plates and MRS agar media was determined, as described before. A specific dilution was determined for each sample using the serial dilution method, and 20-80 µL of the sample was placed and spread into LB-agar with milk 1% plates to visualize protease activity.

### Pupae development success

The total pupae per fly was determined considering the total number of pupae formed at the end of the experiment (14 days after laying time) and the number of female flies permitted to lay eggs in each vial. All pupae formed were counted and visualized using a stereoscopic microscope to differentiate eclosed and non-eclosed pupae. The percentage of pupae success was determined by considering the number of eclosed pupae from the total pupae formed.

### Fecundity

Three females and three males were placed in food with colorant (to contrast embryos), and they were permitted to lay embryos for 48 hours (total time). Every 24 hours, the embryos were counted, and the flies were transferred to a new vial of food with colorant.

### Starvation resistance

Twenty flies were placed in vials containing 1% agar (Sigma-Aldrich, #A1296) after scoring fecundity in each group. Flies were maintained at 25°C and 60-80 % relative humidity. Survival was monitored every three or four hours daily for approximately three days.

### Gene expression

Quantitative RT-PCR analysis was performed to monitor gene expression. Total RNA was purified from 10-15 females (whole-body) using the TRIzol reagent (Invitrogen, #15596018). Complementary DNA (cDNA) was made from 1µg of total RNA with a previous DNase treatment with the DNAse I, Amplification Grade Kit (Invitrogen, #18068015), followed by the iScript cDNA Synthesis Kit (Biorad, #1708891). Quantitative RT-PCR analysis was performed using the SsoAdvanced Universal SYBR Green Supermix kit (Biorad, #1725271) and the CFX96 Real-Time PCR detection system (Biorad). Primer sequences are listed in S1 Table. The data was analyzed using the delta-delta CT method.

## Acknowledgments

We would like to thank all the Broderick Lab members for providing valuable comments that fostered discussion during the project. Authors acknowledge specially to Amber Chou and Natania Kurien for technical assistance during experiments. Krystal Maya Maldonado, Ph.D. is a Latin American Fellow in the Biomedical Sciences, supported by The Pew Charitable Trusts.

## Supporting information captions

**S1 Fig Males dose response to *Pseudomonas entomophila* (*Pe*) infection.** Females and males were used during the experimental strategy for the parental infection using *Pseudomonas entomophila* (*Pe*). As occurs in females, the survival curve shows a dose-response to *Pe* infection in males. Statistical analyses were performed using data over three biological replicates. Log-rank (Mantel-Cox) test was used, and significance is expressed as follows: ns, not significant (p > 0.05); ****p < 0.0001.

**S2 Fig Bacterial load in food and adults. A)** Experimental strategy to study parental infection effects across generations during *Pseudomonas entomophila* (*Pe*) infection. Parents (females and males) were orally infected using three different doses: *Pe* OD100 (High), *Pe* OD50 (Medium), and *Pe* OD5 (Low). Seven days post-infection (dpi), all the parental survivors were collected and randomly transferred to new vials with a density of 25 females and eight males per vial. Flies were permitted to lay eggs for 48 hours to collect the first generation (F1). After laying time, the parents were discarded. A food sample was taken at the start (S) of the experiment, considered as day 0 (days after 48 h of laying time for the parents), and at the end (E) of the experiment, considered as day 10 (10 days post laying time) to determine bacterial load. Once adults appeared, females and males were randomly transferred to new vials to a density of 25 females and eight males per vial, and they were permitted to lay eggs for 48 hours to collect F2 offspring. A food sample was taken at the start (S) and end (E) of the experiment as in the F1 generation. **B-E)** LAB and AAB loads in food and F1 adults. **F-I)** LAB and AAB loads in food and F2 adults. Box plots indicate the median and black points represent one fly per point; whiskers go to the minimum or maximum. Significance is expressed as follows: ns, not significant (p > 0.05); ****p =< 0.0001; **p < 0.005 and, *p < 0.05. Mann Whitney test was used to compare values between the S (start) and E (end) of the experiment, and significant values are shown in the graph. Kruskal-Wallis test, followed by Dunn’s multiple comparisons test, was conducted to compare values according to the parental treatment in each generation per time. Significant values are indicated next to (S) or (E). Treatments labeled with different letters are significantly different by Dunn’s multiple comparisons test, with upper-case letters for (S) and lower-case letters for (E).

**S3 Fig Host physiology strategy, fecundity, and starvation resistance curves. A)** Experimental strategy in detail to study fitness consequences during chronic exposure to the pathogen across generations. The parental infection was conducted as in the previous experiments. Briefly, oral infection in the parents was done using three different doses: *Pe* OD100 (High), *Pe* OD50 (Medium), and *Pe* OD5 (Low). Seven days post-infection (dpi), all the parental survivors were collected and randomly transferred to new vials with a density of 25 females and eight males per vial. Flies were permitted to lay eggs for 48 hours to collect the first generation (F1). At this time, some flies were used to determine fecundity, counting the number of embryos per female, using three females and three males per vial in food supplemented with a green colorant (to permit better contrast when embryos were counted). After laying time, the parents were transferred to 1% agar vials to score starvation resistance. Vials were maintained at 25°C to permit development to the next generation, and total pupae counting was determined at 14 days after the laying time. F1 adults were randomly transferred to new vials, as indicated before, to collect eggs for F2 generation. As in parents, some flies were used to determine fecundity and transferred to food with a green colorant. After laying time, the flies were transferred to 1% agar vials to score starvation resistance. As in F1, vials were maintained at 25°C, permitting development to the next generation, and total pupae counting for F2 was determined at 14 days after the laying time. Once F2 adult emerged, flies were randomly transferred to new vials, and fecundity and starvation resistance were scored as indicated above. Graphs showing total embryos per female are indicated in **B)** for parents, **D)** for F1, and **F)** for F2. The median is indicated per each treatment. Survival curves were obtained for starvation resistance in each generation: **C)** for parents, **E)** for F1, and **G)** for F2. Statistical analyses were performed using the Kruskal-Wallis test, followed by Dunn’s multiple comparisons test for fecundity and the Log-rank (Mantel-Cox) test for survival. Significance is expressed as follows: ns, not significant (p > 0.05); ****p =< 0.0001; ***p < 0.001; **p < 0.005 and, *p < 0.05. Values labeled with different letters are significantly different in Dunn’s multiple comparisons test.

**S4 Fig Correlation between fecundity and starvation resistance.** Median survival in hours for starvation resistance and median of total embryos per female was used to determine the linear correlation between starvation resistance and fecundity. Pearson coefficient for linear association was calculated, and each dot in the graph indicates data from each parental treatment, purple line indicates the linear correlation. Pearson correlation coefficient (r value), 95% confidence interval (CI), R squared (r^2^), and p value are indicated in each graph, **A)** in the parents, **B)** F1, and **C)** F2 offspring.

**Fig S5 Experimental strategy to study the offspring response to new *Pe* infection.** Parental infection and offspring collection was conducted as mentioned before in previous experimental strategies descriptions. Adults from F1 and F2 were newly challenged with Pe using a high dose (*Pe* OD100) at 4 days old. Survival was scored for 7 days and samples for bacterial load were taken at 4, 24 hours and 7 days post-infection. Samples for gene expression were taken at 4- and 24-hours post-infection.

**S6 Fig Bacterial load in offspring after new *Pe* infection. A-D)** Pathogen load is similar between newly *Pe*-challenged groups at 4 hpi and 7 dpi. **E)** Persistence of infection in the unchallenged groups (controls only fed with sucrose solution) varies over time in the first generation. **F, J)** The abundance of microbiome members is similarly independent of the parental treatment when newly challenged with *Pe* at 4 hpi in both generations. **G, K)** Increased abundance of the microbiome is maintained at 7 dpi only in *Pe* OD5 offspring. **H, L)** Microbiome ratio is similar between groups at 4 hpi in both generations. **I)** Microbiome ratio is remodeled only in the control condition at 7 dpi in F1 offspring. **M)** Microbiome ratio is re-established to a similar level between groups at 7 dpi. Box plots indicate the median and black points represent one fly per data point, whiskers go to the minimum or maximum. Mann Whitney test was used to compare bacterial load between pairs unchallenged vs new *Pe* challenged in each parental treatment. For pathogen load, Kruskal-Wallis followed by Dunn’s multiple comparisons test was conducted to compare values between parental treatment in each offspring treatment (between unchallenged only or between new *Pe* challenged only). Significance is expressed as follows: ns, not significant and, *p < 0.05.

**S7 Fig Gene expression in F1 and F2 offspring when newly challenged with *Pe*.** Graphs show mRNA fold change in the unchallenged and newly *Pe* challenged at 4 and 24 hpi of the next genes divided into the following categories **A)** Antimicrobial peptides (AMPs): *Diptericin A* (DptA), *Attacin A* (attA), *Metchnikowin* (Mtk), *Drosomycin-like 3* (Dro3). **B)** Stress response: *Dual oxidase* (Duox), *Glutathione S transferase D5* (GstD5), and *Growth arrest and DNA damage-inducible 45* (Gadd45). **C)** Signaling: *unpaired 3* (upd3), *Suppressor of cytokine signaling at 36E* (Socs36E), and *puckered* (puc). Expression is normalized to the control treatment (unchallenged condition in the parents) for each time and for each offspring treatment (value 1). Bars represent mean ± SD of three biological experiments, each indicating the fold change value obtained in each experiment. Unpaired T-test with Welch correction was used to determine significant differences between the control treatment in the parents and the dose used in each parental treatment following or not following a new *Pe* infection in offspring. Significance is expressed as follows: **p < 0.005 and, *p < 0.05.

**S1 Table Statistical values and primer sequences.**

## References

1. Armitage SA, Genersch E, McMahon DP, Rafaluk-Mohr C, Rolff J. Tripartite interactions: how immunity, microbiota and pathogens interact and affect pathogen virulence evolution. Curr Opin Insect Sci. 2022 Apr;50:100871.

2. Buchmann K. Evolution of Innate Immunity: Clues from Invertebrates via Fish to Mammals. Front Immunol 2014 Sep 23;5:459.

3. Palm NW, De Zoete MR, Flavell RA. Immune–microbiota interactions in health and disease. Clin Immunol. 2015 Aug;159(2):122–7.

4. Acevedo TS, Fricker GP, Garcia JR, Alcaide T, Berasategui A, Stoy KS, et al. The Importance of Environmentally Acquired Bacterial Symbionts for the Squash Bug (*Anasa tristis*), a Significant Agricultural Pest. Front Microbiol. 2021 Oct 4;12:719112.

5. Koga R, Meng XY, Tsuchida T, Fukatsu T. Cellular mechanism for selective vertical transmission of an obligate insect symbiont at the bacteriocyte–embryo interface. Proc Natl Acad Sci. 2012 May 15;109(20):E1230–7.

6. Longdon B, Day JP, Schulz N, Leftwich PT, De Jong MA, Breuker CJ, et al. Vertically transmitted rhabdoviruses are found across three insect families and have dynamic interactions with their hosts. Proc R Soc B Biol Sci. 2017 Jan 25;284(1847):20162381.

7. Lanz-Mendoza H, Contreras-Garduño J. Innate immune memory in invertebrates: Concept and potential mechanisms. Dev Comp Immunol. 2022 Feb;127:104285.

8. Bozler J, Kacsoh BZ, Bosco G. Maternal Priming of Offspring Immune System in *Drosophila*. G3 GenesGenomesGenetics. 2020 Jan 1;10(1):165–75.

9. Tetreau G, Dhinaut J, Gourbal B, Moret Y. Trans-generational Immune Priming in Invertebrates: Current Knowledge and Future Prospects. Front Immunol. 2019 Aug 14;10:1938.

10. Vilcinskas A. Mechanisms of transgenerational immune priming in insects. Dev Comp Immunol. 2021 Nov;124:104205.

11. Broderick NA, Buchon N, Lemaitre B. Microbiota-Induced Changes in Drosophila melanogaster Host Gene Expression and Gut Morphology. McFall-Ngai MJ, editor. mBio. 2014 Jul;5(3):e01117–14.

12. Lesperance DN, Broderick NA. Microbiomes as modulators of *Drosophila melanogaster* homeostasis and disease. Curr Opin Insect Sci. 2020 Jun;39:84–90.

13. Schmidt K, Engel P. Mechanisms underlying gut microbiota–host interactions in insects. J Exp Biol. 2021 Jan 15;224(2):jeb207696.

14. Delbare SYN, Ahmed-Braimah YH, Wolfner MF, Clark AG. Interactions between the microbiome and mating influence the female’s transcriptional profile in *Drosophila melanogaster*. Sci Rep. 2020 Oct 23;10(1):18168.

15. Dobson AJ, Chaston JM, Douglas AE. The *Drosophila* transcriptional network is structured by microbiota. BMC Genomics. 2016 Dec;17(1):975.

16. Freitak D, Wheat CW, Heckel DG, Vogel H. Immune system responses and fitness costs associated with consumption of bacteria in larvae of Trichoplusia ni. BMC Biol. 2007 Dec;5(1):56.

17. Ye YH, Chenoweth SF, McGraw EA. Effective but Costly, Evolved Mechanisms of Defense against a Virulent Opportunistic Pathogen in Drosophila melanogaster. Schneider DS, editor. PLoS Pathog. 2009 Apr 17;5(4):e1000385.

18. Schwenke RA, Lazzaro BP, Wolfner MF. Reproduction–Immunity Trade-Offs in Insects. Annu Rev Entomol. 2016 Mar 11;61(1):239–56.

19. Prakash A, Agashe D, Khan I. The costs and benefits of basal infection resistance vs immune priming responses in an insect. Dev Comp Immunol. 2022 Jan;126:104261.

20. Shianiou G, Teloni S, Apidianakis Y. Intestinal Immune Deficiency and Juvenile Hormone Signaling Mediate a Metabolic Trade-off in Adult Drosophila Females. Metabolites. 2023 Feb 24;13(3):340.

21. Fridmann-Sirkis Y, Stern S, Elgart M, Galili M, Zeisel A, Shental N, et al. Delayed development induced by toxicity to the host can be inherited by a bacterial-dependent, transgenerational effect. Front Genet. 2014 Feb 25;5:27.

22. Morimoto J, Simpson SJ, Ponton F. Direct and trans-generational effects of male and female gut microbiota in *Drosophila melanogaster*. Biol Lett. 2017 Jul;13(7):20160966.

23. Broderick NA, Lemaitre B. Gut-associated microbes of *Drosophila melanogaster*. Gut Microbes. 2012 Jul 14;3(4):307–21.

24. Markow TA, O’Grady P. Reproductive ecology of *Drosophila*. Funct Ecol. 2008 Oct;22(5):747–59.

25. Reaume CJ, Sokolowski MB. The nature of *Drosophila melanogaster*. Curr Biol. 2006 Aug;16(16):R623–8.

26. Liehl P, Blight M, Vodovar N, Boccard F, Lemaitre B. Prevalence of Local Immune Response against Oral Infection in a Drosophila/Pseudomonas Infection Model. Schneider D, editor. PLoS Pathog. 2006 Jun 9;2(6):e56.

27. Moreno-García M, Condé R, Bello-Bedoy R, Lanz-Mendoza H. The damage threshold hypothesis and the immune strategies of insects. Infect Genet Evol. 2014 Jun;24:25–33.

28. Chambers MC, Jacobson E, Khalil S, Lazzaro BP. Consequences of chronic bacterial infection in *Drosophila melanogaster*. Söderhäll K, editor. PLOS ONE. 2019 Oct 24;14(10):e0224440.

29. Vodovar N, Vinals M, Liehl P, Basset A, Degrouard J, Spellman P, et al. *Drosophila* host defense after oral infection by an entomopathogenic *Pseudomonas* species. Proc Natl Acad Sci. 2005 Aug 9;102(32):11414–9.

30. Adamo SA. How insects protect themselves against combined starvation and pathogen challenges, and the implications for reductionism. Comp Biochem Physiol B Biochem Mol Biol. 2021 Aug;255:110564.

31. Mirth CK, Nogueira Alves A, Piper MD. Turning food into eggs: insights from nutritional biology and developmental physiology of Drosophila. Curr Opin Insect Sci. 2019 Feb;31:49– 57.

32. Piper MDW, Zanco B, Sgrò CM, Adler MI, Mirth CK, Bonduriansky R. Dietary restriction and lifespan: adaptive reallocation or somatic sacrifice? FEBS J. 2023 Apr;290(7):1725–34.

33. Schulz NKE, Stewart CM, Tate AT. Female investment in terminal reproduction or somatic maintenance depends on infection dose [Preprint]. Evolutionary Biology; 2022 Jul [cited 2023 Jul 26]. Available from: http://biorxiv.org/lookup/doi/10.1101/2022.07.25.501224

34. Ahlawat N, Maggu K, Jigisha, Arun MG, Meena A, Agarwala A, et al. No major cost of evolved survivorship in *Drosophila melanogaster* populations coevolving with *Pseudomonas entomophila*. Proc R Soc B Biol Sci. 2022 May 11;289(1974):20220532.

35. Barron AJ, Lesperance DNA, Doucette J, Calle S, Broderick NA. Microbiome derived acidity protects against microbial invasion in *Drosophila* [Preprint]. Microbiology; 2023 Jan [cited 2023 Jul 24]. Available from: http://biorxiv.org/lookup/doi/10.1101/2023.01.12.523836

36. Nguyen B, Than A, Dinh H, Morimoto J, Ponton F. Parental Microbiota Modulates Offspring Development, Body Mass and Fecundity in a Polyphagous Fruit Fly. Microorganisms. 2020 Aug 24;8(9):1289.

37. Paniagua Voirol LR, Weinhold A, Johnston PR, Fatouros NE, Hilker M. Legacy of a Butterfly’s Parental Microbiome in Offspring Performance. Johnson KN, editor. Appl Environ Microbiol. 2020 Jun 2;86(12):e00596–20.

38. Schwab DB, Riggs HE, Newton ILG, Moczek AP. Developmental and Ecological Benefits of the Maternally Transmitted Microbiota in a Dung Beetle. Am Nat. 2016 Dec;188(6):679–92.

39. Blum JE, Fischer CN, Miles J, Handelsman J. Frequent Replenishment Sustains the Beneficial Microbiome of *Drosophila melanogaster.* Casadevall A, editor. mBio. 2013 Dec 31;4(6):e00860–13.

40. Lesperance DNA, Broderick NA. Gut Bacteria Mediate Nutrient Availability in *Drosophila* Diets. Appl Environ Microbiol. 2021;87(1).

41. Consuegra J, Grenier T, Baa-Puyoulet P, Rahioui I, Akherraz H, Gervais H, et al. *Drosophila*-associated bacteria differentially shape the nutritional requirements of their host during juvenile growth. PLOS Biol. 2020 Mar 20;18(3):e3000681.

42. Brown LD, Shapiro LLM, Thompson GA, Estévez-Lao TY, Hillyer JF. Transstadial immune activation in a mosquito: Adults that emerge from infected larvae have stronger antibacterial activity in their hemocoel yet increased susceptibility to malaria infection. Ecol Evol. 2019 May;9(10):6082–95.

43. Yap GS. Avirulence: an essential feature of the parasitic lifestyle. Trends Parasitol. 2022 Dec;38(12):1028–30.

44. Fuse N, Okamori C, Okaji R, Tang C, Hirai K, Kurata S. Transcriptome features of innate immune memory in Drosophila. Wappner P, editor. PLOS Genet. 2022 Oct 17;18(10):e1010005.

45. Tang C, Kurata S, Fuse N. Genetic dissection of innate immune memory in *Drosophila melanogaster*. Front Immunol. 2022 Aug 4;13:857707.

46. Radhika R, Lazzaro BP. No evidence for trans-generational immune priming in *Drosophila melanogaster*. Söderhäll K, editor. PLOS ONE. 2023 Jul 13;18(7):e0288342.

47. Wukitch AM, Lawrence MM, Satriale FP, Patel A, Ginder GM, Van Beek EJ, et al. Impact of Chronic Infection on Resistance and Tolerance to Secondary Infection in *Drosophila melanogaster*. Kline KA, editor. Infect Immun. 2023 Mar 15;91(3):e00360–22.

48. Patrnogic J, Castillo JC, Shokal U, Yadav S, Kenney E, Heryanto C, et al. Pre-exposure to non-pathogenic bacteria does not protect *Drosophila* against the entomopathogenic bacterium Photorhabdus. Skoulakis EMC, editor. PLOS ONE. 2018 Oct 31;13(10):e0205256.

49. Acuña Hidalgo B, Armitage SAO. Host Resistance to Bacterial Infection Varies Over Time, but Is Not Affected by a Previous Exposure to the Same Pathogen. Front Physiol. 2022 Mar 21;13:860875.

